# The role of BMP6 in the proliferation and differentiation of chicken cartilage cells

**DOI:** 10.1101/413419

**Authors:** Fei Ye, Hengyong Xu, Huadong Yin, Xiaoling Zhao, Diyan Li, Qing Zhu, Yan Wang

## Abstract

Previous studies have indicated that bone morphogenetic protein (BMP) 6 plays an important role in skeletal system development and progression. However, the mechanism underlying the effects of BMP6 in cartilage cell proliferation and differentiation remains unknown. In this study, cartilage cells were isolated from shanks of chicken embryos and treated with different concentrations of GH. Cell proliferation and differentiation potential was assessed using real-time polymerase chain reaction (RT-PCR) and CCK-8 assays in vitro. The results showed that at 48 h, the Collagen II and BMP6 expression levels in 50 ng/μl GH-treated cartilage cells were significantly higher than in groups treated with 100 ng/μl or 200 ng/μl GH. We further observed that knockdown of BMP6 in cartilage cells led to significantly decreased expression levels of Collagen II and Collagen X. Moreover, the suppression of BMP6 expression by a specific siRNA vector led to significantly decreased expression levels of IGF1R, JAK, PKC, PTH, IHH and PTHrP. Taken together, our data suggest that BMP6 may play a critical role in chicken cartilage cell proliferation and differentiation through the regulation of IGF1, JAK2, PKC, PTH, and Ihh-PTHrP signaling pathways.

## Introduction

Bone morphogenetic proteins (BMPs) are secreted-type multifunctional proteins belonging to the transforming growth factor (TGF)-β superfamily. Many studies have reported that BMPs play very important roles in bone formation and cartilage induction in both vertebrates and invertebrates [1, 2]; moreover, they are also considered crucial molecules involved in cell growth, differentiation, chemotaxis and apoptosis during embryonic development and postnatal tissue remodeling [3]. BMPs stimulate target cells mainly through their specific type I and type II receptors on the cell membrane. When signal transduction occurs, BMPs usually combine with the type II receptor, then increase the expression and activation of receptor type I [4, 5]. BMPs first bind to the receptors on the membrane and transmit this signal through the Smads pathway to promote the differentiation of chondrocytes into the osteogenic lineage [6]. In addition to the Smads signaling pathway, other signaling pathways can also transmit signals from the BMP family, such as mitogen-activated protein kinase (MAPK) pathways [7, 8].

In the BMP family, BMP2, 4 and 6 are all thought to play the most important roles in skeletogenesis. Many studies have suggested that BMP2 is a pivotal signal for the regulation of osteoblastogenesis [9]. Mas et al [10] also showed that BMP2 promotes the expression of Ihh in anterior hypertrophic chondrocytes and the proliferation of chondrocytes. BMP6 is mainly expressed in cartilaginous tissue, where it stimulates mesenchymal cell differentiation into chondrocytes and promote the synthesis of chondrocytes and articular cartilage-specific glycoproteins [11]. BMP6 can also induce the differentiation of MSCs into chondrocytes [12]. In BMPs, BMP6 is a strong factor for bone induction [13]. In addition to the differentiation of MSCs, chondrocytes can be derived from BMSCs, ADSCs and other stem cells induced by BMP6 [14-16]. These findings indicate that BMP6 is an important regulator of bone and cartilage cell proliferation and differentiation. However, the biological activity of BMP6 in cartilage cell proliferation and differentiation, as well as related signaling pathways, has remained unclear. Therefore, a further understanding of the molecular mechanism of BMP6 in cartilage is urgently needed.

In this study, we first extracted and cultured cartilage cells from different breeds of chickens, and we then investigated the expression of BMP6 and the changes in expression of key genes involved in related signaling pathways through GH-mediated induction at different concentrations to determine its potential role in cell proliferation and differentiation. Finally, to explore the mechanism of BMP6-mediated effects on the proliferation and differentiation of cartilage cells, we modulated the expression of BMP6 through siRNA and measured its effects by quantitative real-time PCR analysis. Collectively, this study provides evidence that within cartilage cells, BMP signaling regulates genes associated with both cell proliferation and differentiation.

## Materials and Methods

### Animals

Avian broiler and Yellow bantam chickens, which have major differences, were used in this study. Avian broilers were provided by the Zheng Da Company (Chengdu, China). Yellow bantam chickens were provided by the Jin Ling Company (Guangzhou, China). All animal studies were performed in accordance with appropriate guidelines. All experimental protocols were approved by the Committee on Experimental Animal Management of Sichuan Agricultural University, permit number 2014-18.

### Cell culture

The eggs were incubated for 15 days after sterilization. Primary cartilage cells were isolated from the shank of 15-day-old chicken embryos with 0.25% trypsin (Gibco, USA), digestion for 0.5 h and then 0.1% collagenase Ⅱ (Gibco, USA) for 1.5 h at 37°C under sterile conditions. The cells were grown in DMEM/F12 medium (Gibco, USA) supplemented with 10% fetal bovine serum (FBS, Gibco, USA), 100 U/ml penicillin and 100 U/ml streptomycin (Gibco, USA) in a humidified atmosphere of 5% CO_2_ at 37°C.

### Immunofluorescence

Cartilage cells were fixed in 4% paraformaldehyde for 20 min after adhering for 24 h in the incubator and were then washed three times in PBS, 5 min per wash. The cells were permeabilized with 0.5% Triton-X-100 (Gibco, USA) for 15 min at room temperature and then washed three times in PBS, 5 min per wash. The cells were blocked with Blocking buffer (Bio-Rad, USA) for 2 h at 37°C and then washed as above. The cells were incubated in PBS containing an antibody towards type II collagen (1:1000) (Abcam, USA) overnight at 4°C. The following day, the cells were washed as described above and incubated in PBS containing IgG (1:250) (Abcam, USA) for 2 h at 37°C under dark conditions. After washing three times, the cells were counterstained with DAPI. Photomicrographs were taken using an Olympus digital camera system.

### Cell viability assay

Cell proliferation activity was evaluated via Cell Counting Kit-8 assay (CCK8, Bioss, China). At different time points, the medium was replaced with 100 μl of fresh medium containing 10 μl of CCK8. Three hours after the addition of CCK8, cell viability was determined with a microplate reader (Thermo Electron, USA) at a wavelength of 450 nm. All plates had control wells containing medium without cells to obtain a value for background luminescence, which was subtracted from the test sample readings. Each experiment was performed in triplicate.

### Cell treatments

Cells were treated with 0 ng/μl, 50 ng/μl, 100 ng/μl, and 200 ng/μl GH while cells were grown to 80% confluence. Cells were harvested after transfection for 24 h, 48 h, and 72 h. Each experiment was performed in triplicate.

Two siRNAs were designed based on the sequence from NCBI (XM-418956.4) and are listed in Table 1. The cartilage cells were grown to 80% confluence and transfected with 5 μl of Lipofectamine 2000 (Invitrogen, USA) and 5 μl of siRNA (Sangon Biotech, China) according to the manufacturer’s instructions. Cells were harvested after transfection at 24 h, 48 h, and 72 h. Each experiment was performed in triplicate.

Table 1: Sequences of siRNA targeting the BMP6 gene

### RNA Extraction and qRT-PCR

Total RNA was extracted from cells using the RNAiso plus Reagent (Takara, Japan) according to the manufacturer’s instructions. The first-strand cDNA was synthesized using the PrimeScript™ RT reagent Kit reverse transcriptase (Takara, Japan) with 1 μg of total RNA according to the manufacturer’s instructions. β-actin was chosen as an internal standard to control for variability in amplification due to differences in the starting mRNA concentrations. The forward and reverse primer sequences used to amplify the genes were designed according to the sequence retrieved from the NCBI and listed in Table 2. Quantitative PCR was performed using SYBR Green PCR technology with a Bio-Rad CFX Connect Real Time System (Bio-Rad, USA). PCR was performed at 98°C for 120 s, followed by 40 amplification cycles (98°C for 2 s, X°C for 15 s, 72°C for 10 s), followed by 65°C to 95°C per second and then 4°C forever. The relative gene expression level was calculated using the 2^-ΔΔC_T_^ method. All PCR runs were performed in triplicate.

Table 2: Primer sequences for qRT-PCR

### Statistical analyses

All cell culture experiments were performed a minimum of three times. Statistical analyses were conducted using SAS 8.0 software for Windows. All data are expressed as means ± SEM, and statistical analysis was performed using Student’s t-test. P-values < 0.05 were considered statistically significant.

## Results

### Immunofluorescence of cartilage cells

Collagen II as a special marker for cartilage cells was imaged in cartilage cells isolated from avian broiler and yellow bantam chickens, and the cells morphologies of the two breeds was consistent (Fig 1). These results indicate that the cells we cultured were cartilage cells.

**Fig 1:**
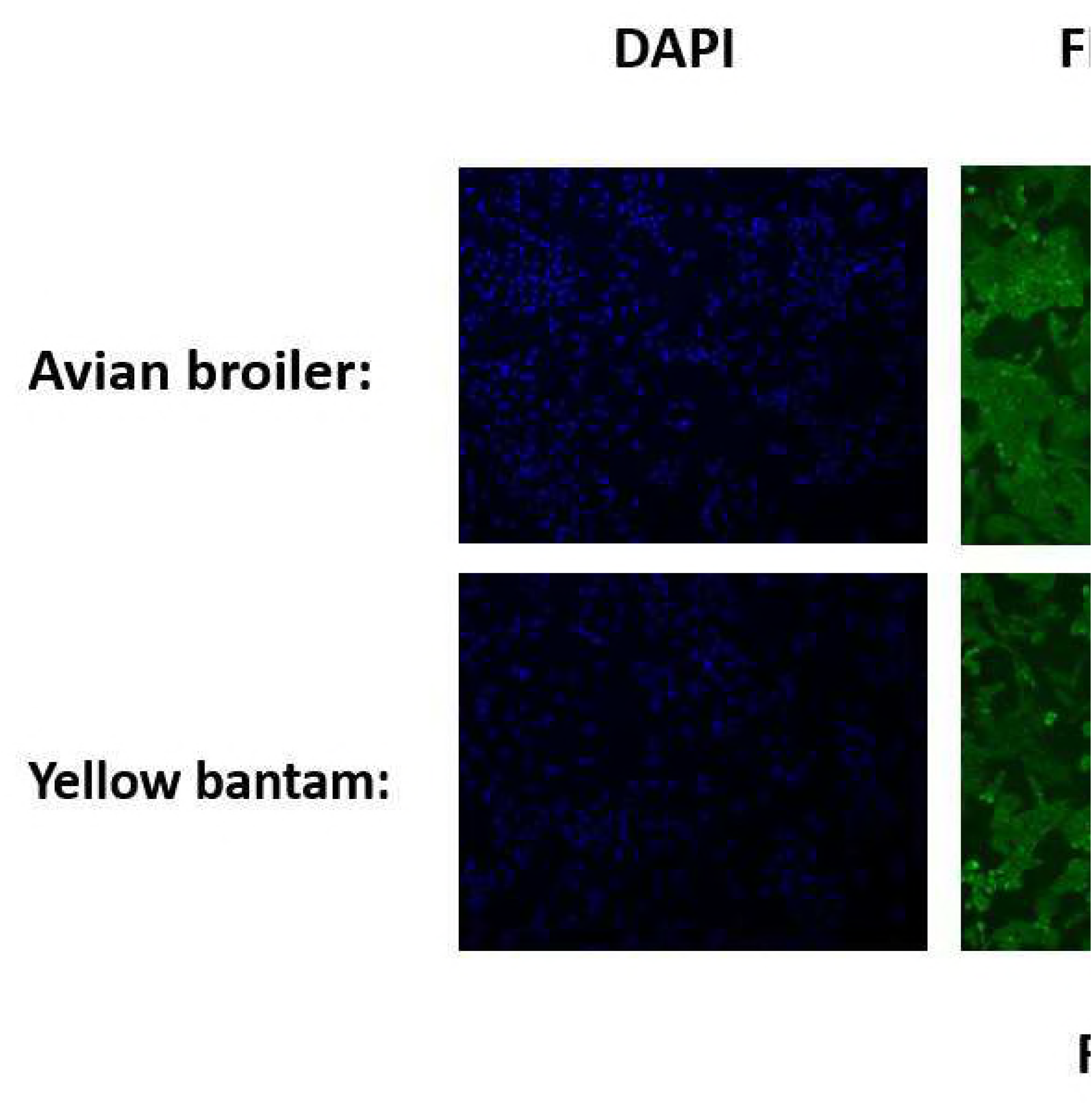
Immunofluorescence of markers in cartilage cells. Nuclei stained with DAPI are shown in the left panels. The pictures above indicated that staining of the cells for the marker collagen II was positive. The merged images are shown in the right-most panels. Scale bar = 100 μm.

### Growth Kinetics of cartilage cells of the two breeds

The growth kinetics of the cartilage cells from the two breeds at different timepoints are shown by the growth curves. Avian broiler cartilage cells entered the logarithmic phase after approximately 3 days, which ended at the ninth day, whereas yellow bantam cartilage cells entered the logarithmic after approximately 4 days and ended at the tenth day. Over additional days, the ability of cells to grow decreased (Fig 2).

**Fig 2:**
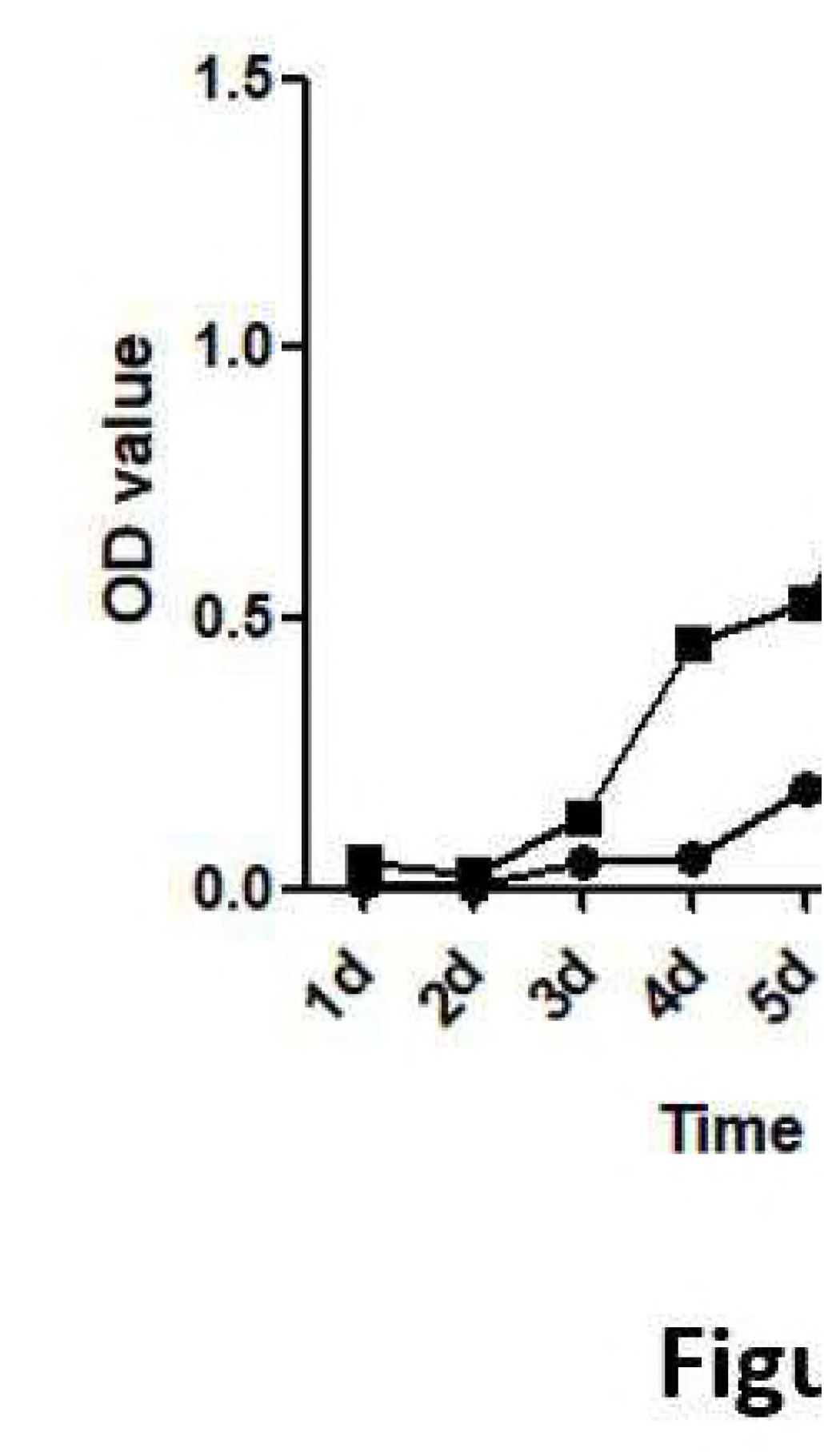
Growth curves of chicken cartilage cells. The growth curves of cells were typically sigmoidal, with cell density reflected by the vertical axis. The growth curve consisted of a latent phase, a logarithmic phase, and a plateau phase (n=3).

### Expression of *BMP6* mRNA in cartilage cells of the two chicken breeds

The relative expression level of BMP6 mRNA was detected in cartilage cells from avian broilers and yellow bantams at day 0, day 1, day 2, day 3, day 4, and day 5 according to the growth curves of cells (Fig 3). Real-time PCR experiments showed that the cellular expression of BMP6 significantly (P<0.05) increased at the fourth day and fifth day, which was consistent with the growth curves.

**Fig 3:**
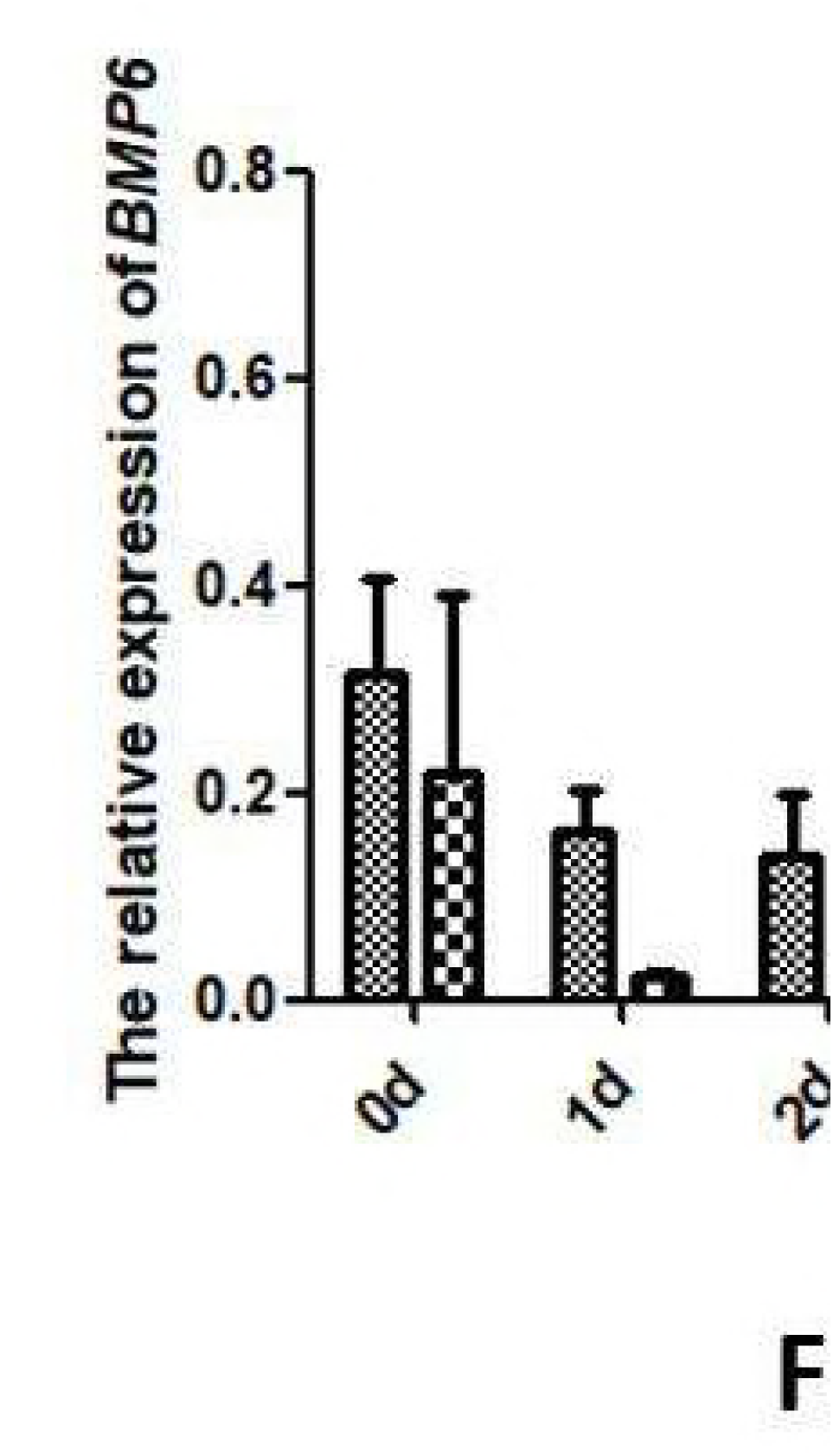
The relative expression level of BMP6 in cartilage cells of avian broilers and yellow bantams at different days. All values are presented as the means ± SEM (n=3). (*) represents statistical significance (P<0.05).

### Expression of *BMP6* mRNA after GH induction

Based on the growth curves and the relative expression of BMP6 mRNA in cartilage cells of avian broilers and yellow bantams, we selected cartilage cells of avian broilers for induction by GH and interference by siRNA targeting BMP6. Collagen II, a special marker for cartilage cells, was detected in cartilage cells after GH-induction. Relative to the β-actin gene, the expression levels of BMP6 mRNA varied considerably at different times (Fig 4). As seen from the results (Fig 4(A)), the proliferation of cartilage cells in the three treatment groups at 24 h and 7 2 h was not significantly higher than that in the blank treatment group (P<0.05). However, at 48 h, the proliferation of cartilage cells was significantly elevated in the treatment group compared to that of the blank group, 100 ng/μl and 200 ng/μl groups (P<0.05). These results indicated that 50 ng/μl GH was the most sensitive to the proliferation induction of cartilage cells. The relative expression levels of BMP6 mRNA in cartilage cells after induction are shown in Fig 4 (B). In the 50 ng/μl- and 100 ng/μl-treatment groups, the expression of BMP6 mRNA was significantly increased relative to the blank treatment group and 200 ng/μl at 48 h, which was consistent with the expression of Collagen II mRNA (P<0.05). This showed that the expression of BMP6 mRNA was significantly increased in the process of proliferation of cartilage cells.

**Fig 4:**
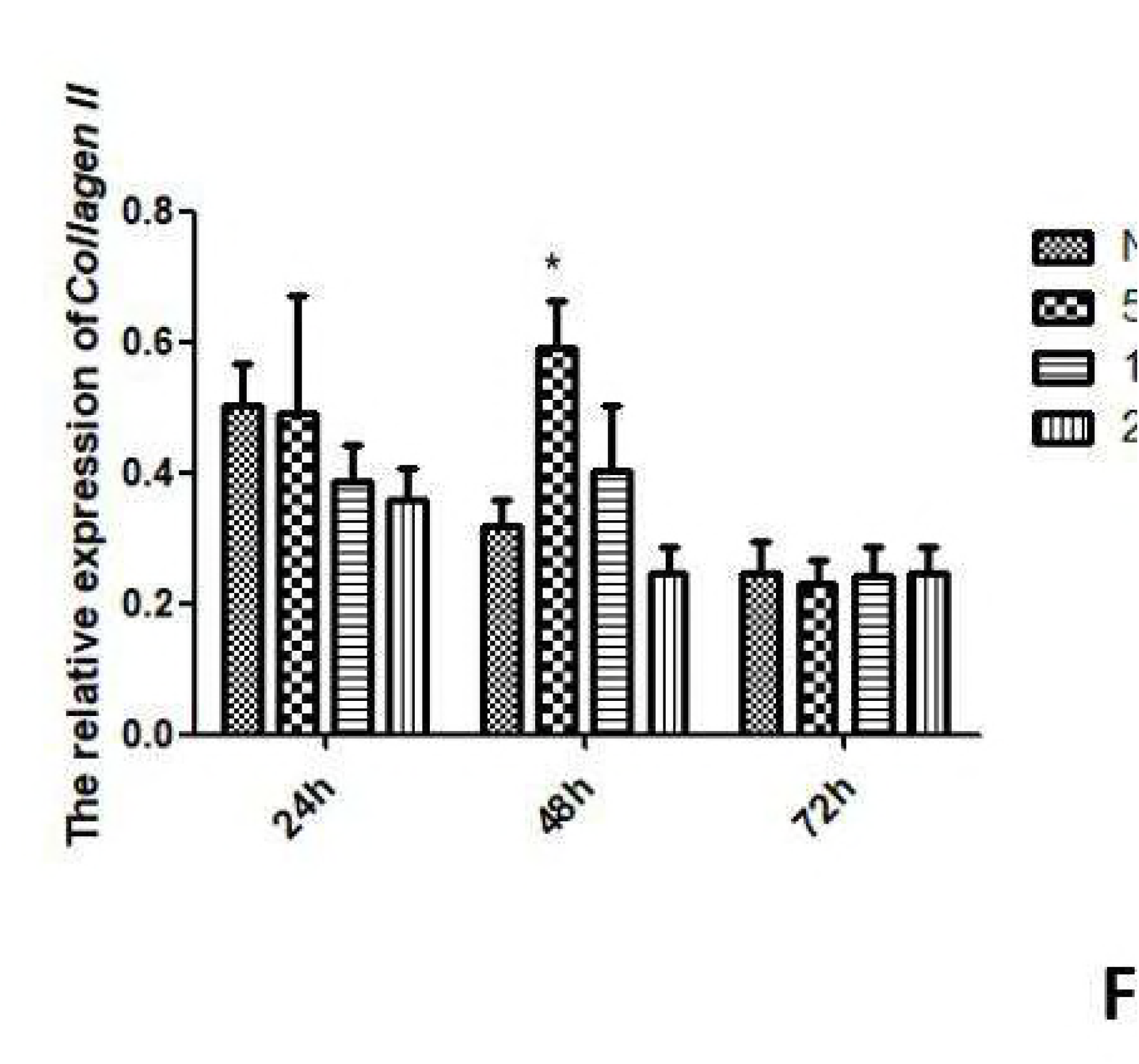
The relative expression of Collagen II (A) and BMP6 (B) mRNA in cartilage cells after GH induction. All values are represented as the means ± SEM (n=3). (*) represent statistical significance (P<0.05).

### The interference efficiency of the two siRNAs

It can be seen from Fig 5 that the relative expression of BMP6 mRNA of cartilage cells after interference with siBMP6.1 and siBMP6.2 at 24 h was not significantly (P>0.05) different from the control group, indicating that the efficiency of siRNA interference was not reflected at this time. At 48 h, the expression levels of BMP6 mRNA in the treatment groups were significantly (P<0.05) decreased, to 25% and 25%, respectively, compared with the control group. At 72 h, the expression levels of BMP6 mRNA in the siRNA groups were significantly (P<0.05) decreased, to 34% and 29%, respectively, compared with the control group. The interference efficiency of the two siRNAs at 48 h and 72 h were marked and stable, indicating their utility for subsequent experiments.

**Fig 5:**
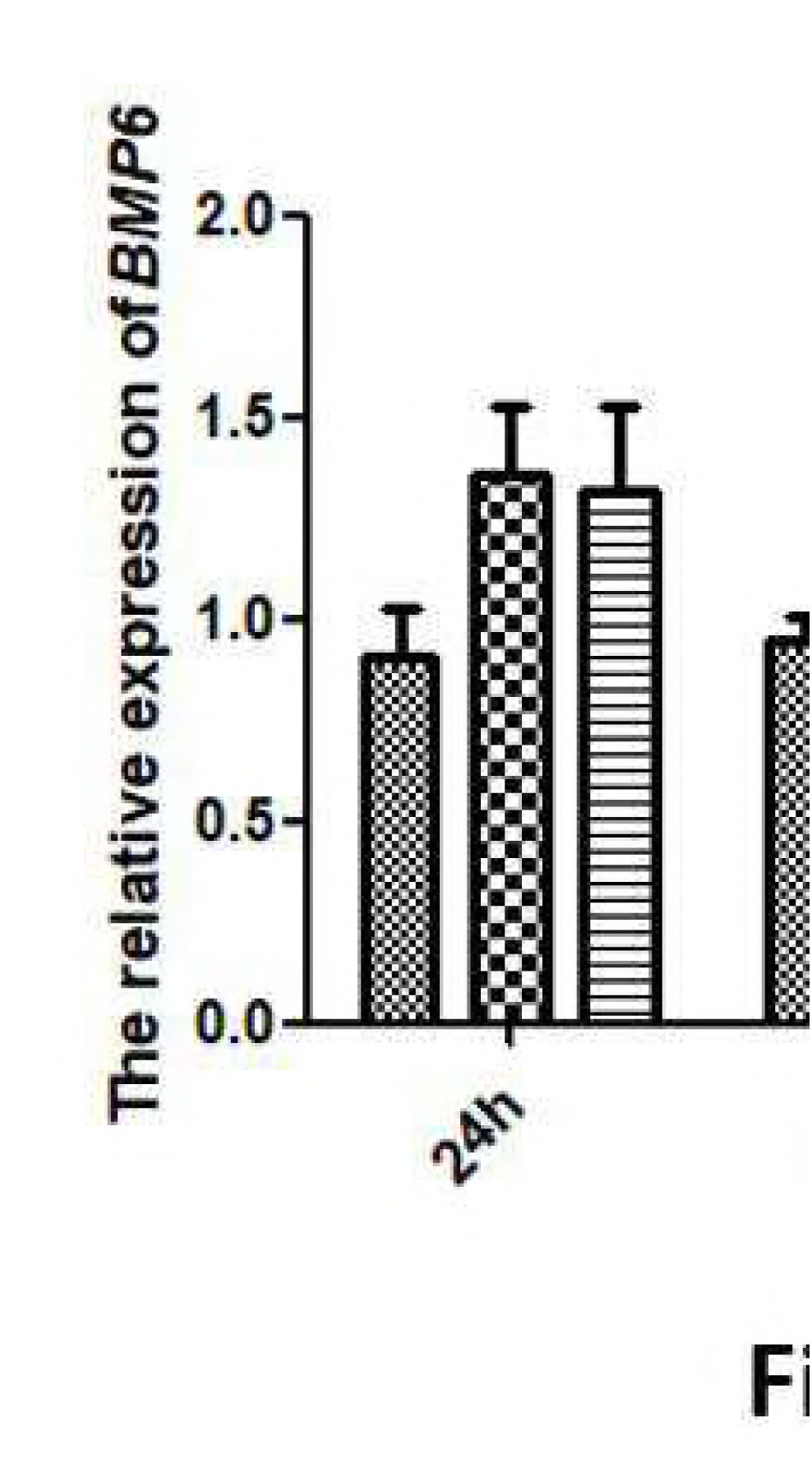
The interference efficiency of the two siRNAs. All values represent means ± SEM (n=3). (*) represents statistical significance (P<0.05).

### The expression levels of *Collagen II* and *Collagen X* mRNA after interference with *BMP6*

Collagen II and Collagen X have been suggested to be important regulators of chondrocyte proliferation and differentiation. We therefore further measured their gene expression levels after interfering with BMP6 expression. As shown in Fig 6, we found that at 48 h and 72 h after interference with BMP6, the Collagen Ⅱ and Collagen X mRNA expression levels in chondrocytes were significantly lower than in the control group (P < 0.05).

**Fig 6:**
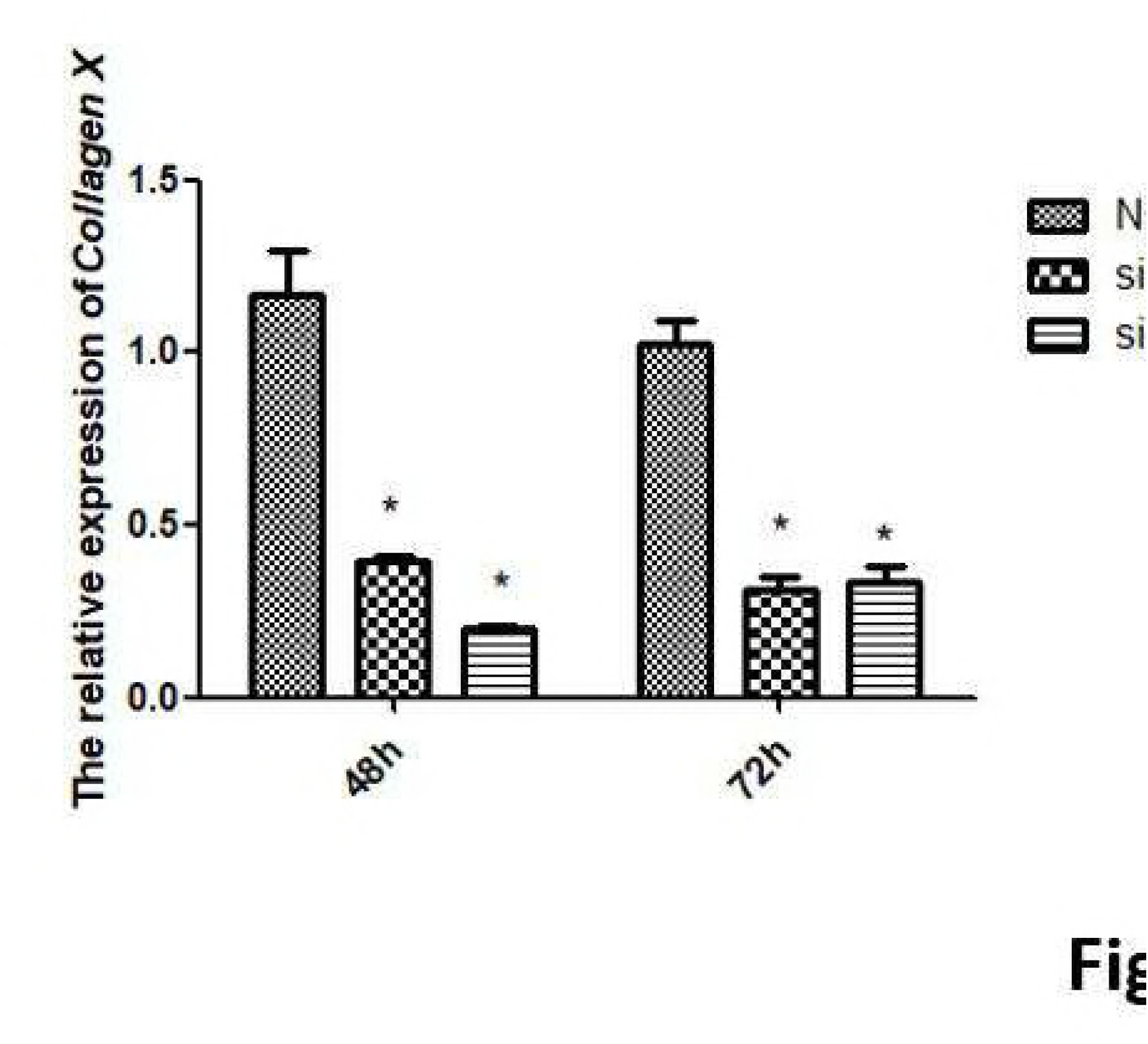
The relative expression levels of Collagen II and Collagen X mRNA in cartilage cells after siRNA treatment. All values are represented as the means ± SEM (n=3). (*) represents statistical significance (P<0.05).

### The expression levels of *IGF1R, JAK, PKC, PTH* and *PTHrP* mRNA after interference with *BMP6*

Since IGF1, PKC, JAK2/STAT, Ihh/PTHrP and PTH signaling have been reported to affect the proliferation and differentiation of chondrocytes, we further investigated whether BMP6 was also involved in these processes. We used siRNA to knock down BMP6 gene expression in plasmid-transfected cells. Real-time PCR analysis showed that transfecting chondrocytes with BMP6-targeting siRNA resulted in a significant inhibition of expression of IGF1R, JAK, PKC, PTH and PTHrP at 72 h (Fig 7). However, at 48 h, the relative expression of IGF1R, JAK, PKC, PTH and PTHrP mRNAs had no significant (P > 0.05) differences between the treatment and control cartilage cells, whereas the relative expression of IHH in the treatment groups was significantly (P < 0.05) lower than in the control group.

**Fig 7:**
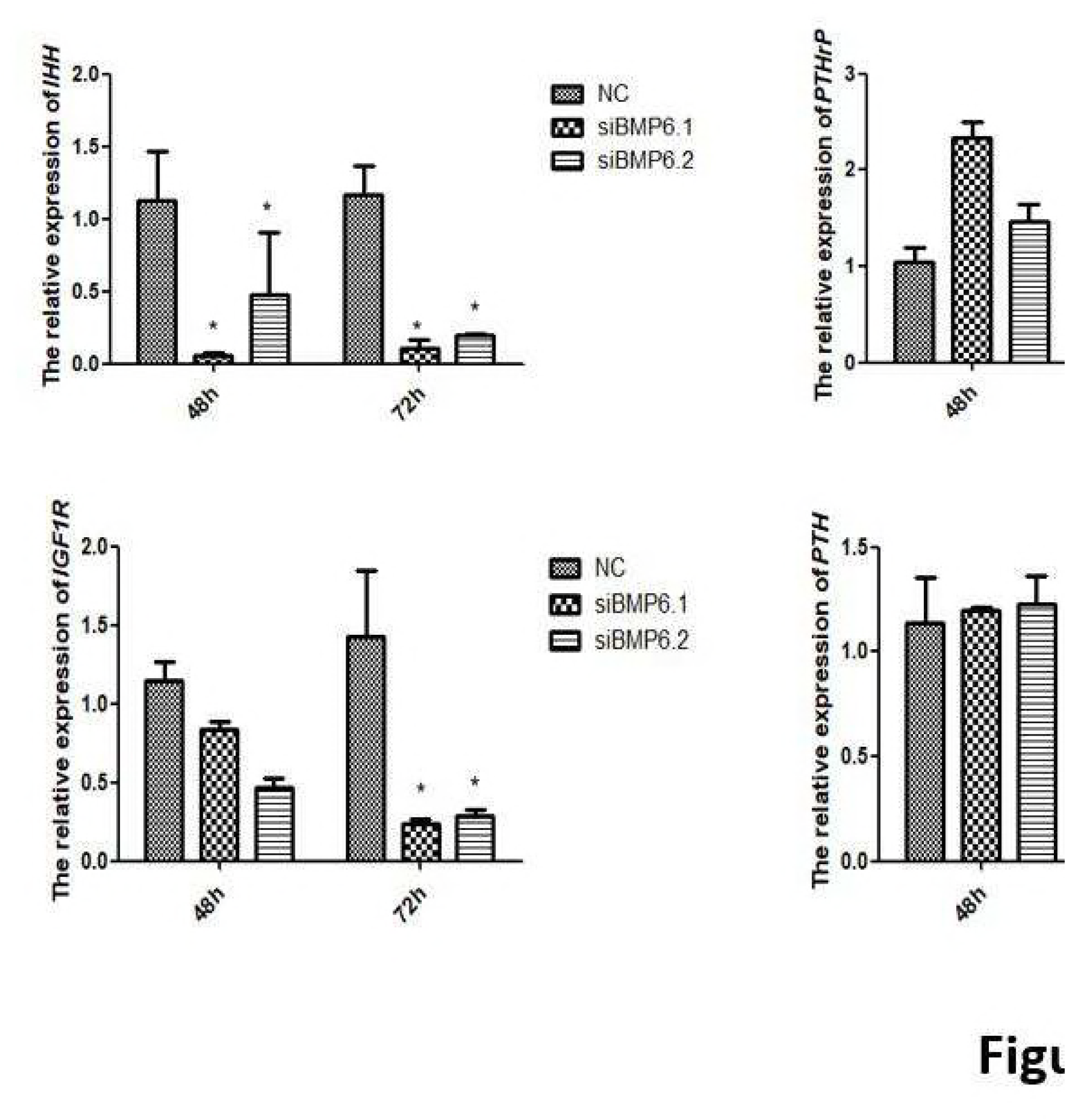
The relative mRNA expression of selected genes in cartilage cells after siRNA treatment. All values represent means ± SEM (n=3). (*) represents statistical significance (P<0.05).

## Discussion

Currently, studies regarding the BMP6 gene in chondrocytes have primarily focused on humans, mice, rabbits and other mammals, with few studies focused on chickens. We used immunofluorescence to identify and observe the cell morphology, used CCK8 assays to detect the proliferation of cartilage cells and real-time PCR to detect the BMP6 mRNA expression in cartilage cells cultured from avian broiler and yellow bantam chickens. Collagen II is a representative protein indicating the proliferation of chondrocytes, and thus can be used as a cell marker; many studies have examined the expression of type II collagen in the mandibular condyle, using both immunohistochemistry and in situ hybridization techniques[17-24]. Our results showed that the avian and yellow bantam cartilage cells had no significant difference. According to these results, we selected cartilage cells cultured from avian chicken growth plates to research the function of BMP6 for regulating the proliferation and differentiation of cartilage cells.

GH is an important regulatory factor for longitudinal growth of the bone [25]. Local injection of GH can increase the number of cartilage cells in rats [26]. The expression of GH can promote the expression of Collagen II mRNA in cartilage cells and maintain a chondrocyte phenotype. We used GH to induce the cartilage cells in order to detect the expression of BMP6 mRNA in proliferative cells. The results showed that the sensitivity of cartilage cells to GH varied with GH concentration. When cartilage cells proliferated, the expression of BMP6 was significantly increased. Therefore, we speculated that BMP6 participates in the proliferation of cartilage cells.

RNA interference uses homologous double-stranded RNA (dsRNA) to induce the silencing of specific target genes and block gene activity. SiRNA (small interfering RNA) is the intermediate of RNA interference, which is necessary for the RNA interference and can stimulate the complementary target mRNA silencing. Collagen X is specifically expressed in hypertrophic chondrocytes [27]. In this experiment, two siRNAs were designed and synthesized according to the BMP6 gene CDS region, and the expression of BMP6 mRNA was inhibited. BMP6 was involved in the proliferation of cartilage cells, and, when the expression of BMP6 mRNA was suppressed in cartilage cells, the expression of Collagen II and Collagen X mRNA—two marker genes representing the proliferation and differentiation of cartilage cells—decreased in cartilage cells. These results showed that the proliferation and differentiation of chondrocytes were both blocked following the disruption of the expression of BMP6. Together with the previous results, we speculate that BMP6 is involved in the proliferation and differentiation of cartilage cells, which is consistent with the existing research findings.

IGF1 plays an important role in the growth and development of bone cells; it can regulate the function of osteoblasts in various forms and participates in bone reconstruction. IGF1 can significantly promote the proliferation of bone progenitor cells both in vivo and in vitro [28]. The signal transmission mediated by IGF1 is specifically induced by the IGF1R on the cell surface, thus promoting the growth, differentiation and apoptosis of tissue cells. To explore whether IGF1R is involved in the regulation of chondrocyte proliferation and differentiation of BMP6, the expression of IGF1R mRNA was detected while using RNAi technology to interfere with the expression of BMP6. When the expression of BMP6 was inhibited, the expression of IGF1R was also inhibited. Therefore, we speculate that BMP6 has a regulatory effect on IGF1.

JAK2 is a common signaling pathway induced by multiple cytokine- and growth factor-mediated signaling within cells [29]. When JAK activity is inhibited, osteoarthritis articular chondrocytes can reproduce and differentiate normally [30]. Interleukin-6 and interleukin-7 can induce cartilage cell activity through the JAK2/STAT signaling pathway [31, 32]. It has been found that BMP7 can promote osteogenic differentiation of osteoblasts through JAK2/STAT5B signaling [33], and we hypothesized that BMP6 may regulate the proliferation and differentiation of chondrocytes through the JAK2/STAT signaling pathway. To explore whether JAK2 is involved in the regulatory effects of BMP6 on the proliferation and differentiation of cartilage cells, we used RNAi technology to interfere with the expression of BMP6 and subsequently detect the expression of JAK2 in the JAK2/STAT signaling pathway. Our results showed that the expression of JAK2 was also inhibited when the expression of BMP6 was decreased. We hypothesize that BMP6 has regulatory effects on JAK2 expression.

PKC is an important substance in cell signal transduction pathways and participates in the process of proliferation and differentiation of chondrocytes; it is one of the important signal transducers affecting the growth and development of cartilage and its eventual degeneration [34]. PKC signaling and multiple other signaling pathways participate in IGF1 induction of chondrocyte proliferation and differentiation [35]. Estrogen and vitamin D-mediated signaling depends on the PKCα pathway to regulate the activity of cartilage cells [36]. To explore whether PKC is involved in the BMP6-mediated regulation of the proliferation and differentiation of cartilage cells, we used RNAi technology to interfere with the expression of BMP6 and detect the subsequent expression of PKC. Our results showed that PKC expression was inhibited when the expression of BMP6 was inhibited. We hypothesize that PKC is involved in the regulation of chondrocyte proliferation and differentiation mediated by BMP6.

The main function of PTH is to regulate Ca2^+^ and Phosphorous metabolism and promote bone absorption. PTH stimulates the expression of Collagen II in cartilage cells [37]. A previous study found that PTH interacts with the TGF-beta signaling pathway at the receptor level [38]. BMP6 belongs to the TGF-beta family, therefore we speculated that PTH may participate in the regulation of chondrocytes by BMP6. We used RNAi technology to interfere with the expression of BMP6 and detected subsequent PTH expression. Our results showed that PTH expression was also inhibited when the expression of BMP6 was inhibited. We hypothesize that PTH participates in the regulation of chondrocyte proliferation and differentiation by BMP6. Ihh/PTHrP signaling is important in bone development regulation, modulating the differentiation of chondrocytes and osteoblasts, maintaining the cartilage cell proliferation state [39] and determining the length of the growth plate cartilage by regulating the bone longitudinal growth rate [40, 41]. PTHrP can inhibit osteoblast differentiation by down-regulating BMP2 expression [41]. We hypothesized that the Ihh/PTHrP signaling pathway may be involved in the regulation by BMP6 of chondrocytes. We used RNAi technology to interfere with the expression of BMP6 and detect the subsequent expression of Ihh and PTHrP. Our results showed that the expression of Ihh and PTHrP mRNA was also inhibited when the expression of BMP6 mRNA was decreased. The Ihh/PTHrP signaling pathway participates in the regulation by BMP6 of the proliferation and differentiation of cartilage cells.

BMP6 belongs to the TGFβ family, and current research indicates that the regulation of chondrocytes is mainly achieved through the Smad signaling pathway of TGFβ [42]. BMP2, BMP9 and BMP6 belong to the BMP family. BMP2 can increase the proliferation of chondrocytes and the elongation of chondrocytes by inducing the expression of Ihh [10]. BMP9 and GH can synergize the osteogenic differentiation of mesenchymal stem cells through the JAK/STAT/IGF1 signaling pathway. As one of the strongest members of the BMPs family, BMP6 may regulate the proliferation and differentiation of chondrocytes through other signaling pathways in addition to the Smad signaling pathway. Therefore, we analyzed the IGF1/JAK/PKC/PTH/Ihh-PTHrP signaling pathways, which play important roles in the process of proliferation and differentiation of cartilage cells, and detected the expression changes of several key genes (IGF1R, JAK2, PKC, PTH, Ihh, PTHrP) in these signaling pathways when the expression of BMP6 mRNA was inhibited. The results collectively showed that the expression of these genes decreased significantly. IGF1/JAK/PKC/PTH/Ihh-PTHrP are involved in the regulation by BMP6 of the proliferation and differentiation of cartilage cells.

## Conclusion

There was no significant difference in cartilage cells cultured from different chicken breeds. BMP6 was highly expressed when cartilage cells proliferated. When the expression of BMP6 mRNA was decreased, the proliferation and differentiation of chondrocytes was blocked, indicating that BMP6 is involved in the proliferation and differentiation of cartilage cells. The expression levels of key genes involved in the IGF1R, JAK2, PKC, PTH, IHH and PTHrP signaling pathways were significantly lower in cartilage cells when the expression of BMP6 mRNA was decreased. These key genes involved in signaling pathways were involved in the regulation by BMP6 of the proliferation and differentiation of cartilage cells.

## Acknowledgments

Conceived and designed the experiments: Y.W. and Q.Z.; Performed the experiments: F.Y., H.Y.X., H.D.Y.; Analyzed the data: F.Y.; Contributed reagents/materials/analysis tools: H.Y.X., D.Y.L., X.L.Z.; Wrote the paper: F.Y.; Revised the manuscript: Y.W. and Q.Z.

## Supporting information

**S1 Table 1: Sequences of siRNA targeting the BMP6 gene**

**S2 Table 2: Primer sequences for qRT-PCR**

**S1 Fig 1: Immunofluorescence of markers in cartilage cells.** Nuclei stained with DAPI are shown in the left panels. The pictures above indicated that staining of the cells for the marker collagen II was positive. The merged images are shown in the right-most panels. Scale bar = 100 μm.

**S2 Fig 2: Growth curves of chicken cartilage cells.** The growth curves of cells were typically sigmoidal, with cell density reflected by the vertical axis. The growth curve consisted of a latent phase, a logarithmic phase, and a plateau phase (n=3).

**S3 Fig 3: The relative expression level of BMP6 in cartilage cells of avian broilers and yellow bantams at different days.** All values are presented as the means ± SEM (n=3). (*) represents statistical significance (P<0.05).

**S4 Fig 4: The relative expression of Collagen II (A) and BMP6 (B) mRNA in cartilage cells after GH induction.** All values are represented as the means ± SEM (n=3). (*) represent statistical significance (P<0.05).

**S5 Fig 5: The interference efficiency of the two siRNAs.** All values represent means ± SEM (n=3). (*) represents statistical significance (P<0.05).

**S6 Fig 6: The relative expression levels of Collagen II and Collagen X mRNA in cartilage cells after siRNA treatment.** All values are represented as the means ± SEM (n=3). (*) represents statistical significance (P<0.05).

**S7 Fig 7: The relative mRNA expression of selected genes in cartilage cells after siRNA treatment.** All values represent means ± SEM (n=3). (*) represents statistical significance (P<0.05).

